# Unbiased population heterozygosity estimates from genome-wide sequence data

**DOI:** 10.1101/2020.12.20.423694

**Authors:** Thomas L Schmidt, Moshe Jasper, Andrew R Weeks, Ary A Hoffmann

## Abstract

1. Heterozygosity is a metric of genetic variability frequently used to inform the management of threatened taxa. Estimating observed and expected heterozygosities from genome-wide sequence data has become increasingly common, and these estimates are often derived directly from genotypes at single nucleotide polymorphism (SNP) markers. While many SNP markers can provide precise estimates of genetic processes, the results of ‘downstream’ analysis with these markers may depend heavily on ‘upstream’ filtering decisions.
2. Here we explore the downstream consequences of sample size, rare allele filtering, missing data thresholds and known population structure on estimates of observed and expected heterozygosity using two reduced-representation sequencing datasets, one from the mosquito *Aedes aegypti* (ddRADseq) and the other from a threatened grasshopper, *Keyacris scurra* (DArTseq).
3. We show that estimates based on polymorphic markers only (i.e. SNP heterozygosity) are always biased by global sample size (N), with smaller N producing larger estimates. By contrast, results are unbiased by sample size when calculations consider monomorphic as well as polymorphic sequence information (i.e. genome-wide or autosomal heterozygosity). SNP heterozygosity is also biased when differentiated populations are analysed together, while autosomal heterozygosity remains unbiased. We also show that when nucleotide sites with missing genotypes are included, observed and expected heterozygosity estimates diverge in proportion to the amount of missing data permitted at each site.
4. We make three recommendations for estimating genome-wide heterozygosity: (i) autosomal heterozygosity should be reported instead of (or in addition to) SNP heterozygosity; (ii) sites with any missing data should be omitted; (iii) populations should be analysed in independent runs. This should facilitate comparisons within and across studies and between observed and expected measures of heterozygosity.

## Introduction

The power provided by single nucleotide polymorphism (SNP) markers detected using genome-wide sequencing approaches is leading to their increased use in conservation genetic studies (Garner et al., 2016). SNPs are popular for investigating levels of genetic differentiation among remnant populations and for comparing levels and patterns of genetic variation within populations (Campbell et al., 2019; Maroso et al., 2016) which provides information on the adaptive potential of populations (Ørsted et al., 2019) as well as patterns of inbreeding and relatedness (Mulvena et al., 2020). Results from SNP studies are being interpreted for use in management decisions that include genetic rescue, genetic mixing and founder selection in threatened species programs (Fitzpatrick et al., 2020). The relative ease of generating SNP genotypes is leading to their increased use by nonspecialists, particularly through the availability of companies such as Diversity Arrays Technology (https://www.diversityarrays.com), which provide SNP genotypes through customised in-house processes (Gruber et al., 2017; Mulvena et al., 2020; Wright et al., 2019).

Considering the popularity of SNP markers, it is important to be aware of any biases inherent in their application to conservation genetics and elsewhere. While potential biases have been considered for the detection of structure between populations (Linck & Battey, 2019; Wright et al., 2019), there has been less focus on the estimation of genetic variability within populations. These estimates are important because they link to the evolutionary potential of populations, which is typically higher in populations with greater genetic variability (Hoffmann et al., 2017; Ørsted et al., 2019). Genetic variability of populations is therefore crucial when making genetic management decisions for threatened species (Hoffmann et al., 2020; Weeks et al., 2011).

Genetic variation in populations is measured in several ways, the most common of which are heterozygosity (observed and expected) and the proportion of nucleotide sites that are polymorphic. Heterozygosity is usually estimated from a substantial number of individuals sampled from each population, but with large quantities of sequence data fewer individuals may be needed (Nazareno et al., 2017). Accurate heterozygosity estimates also require that the apparent diversity at a site is not related to errors introduced during sequencing or genotyping, the latter of which requires adequate coverage to ensure that both strands of a diploid individual are sequenced (Nielsen et al., 2011). While expected heterozygosity is estimated from allele frequencies, observed heterozygosity is estimated from individual genotypes directly and depends on both the amount of genetic variation in the population and the level of inbreeding, which increases homozygosity (Ritland, 1996). Inbreeding can thus be estimated by comparing observed heterozygosity to expected heterozygosity, with the latter expected to be relatively higher when there is inbreeding (F_IS_ > 0).

For heterozygosity, *h_i_*, the observed heterozygosity for an individual at site *i* can be averaged across *n* sites as 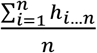 and averaged across a sample of individuals to provide a population estimate. This can be calculated from variation at polymorphic sites only (i.e. SNP heterozygosity) or at both polymorphic and monomorphic sites (i.e. genome-wide or autosomal heterozygosity). Both SNP heterozygosity and autosomal heterozygosity appear in the literature; most population-focussed studies tend to report SNP heterozygosity (Bock et al., 2018; Chen et al., 2016; Jones et al., 2012; Mathur et al., 2019; Surbakti et al., 2020) although others use autosomal heterozygosity (Hohenlohe et al., 2010) which is the parameter reported in studies comparing individual genomes (Gopalakrishnan et al., 2017; Westbury et al., 2019). As SNP heterozygosities will be orders of magnitude larger than autosomal heterozygosities, the two parameters cannot be directly compared, though for studies of a single population autosomal heterozygosity can be converted to SNP heterozygosity by dividing the estimate by the proportion of polymorphic sites. Fig. S1 provides a visualisation of how observed heterozygosity is calculated using all sequence information (autosomal heterozygosity) and using polymorphic markers only (SNP heterozygosity).

This paper investigates how SNP heterozygosity and autosomal heterozygosity perform under variable conditions of sampling and filtering. These include local and global sample size, rare allele filtering, missing data thresholds and the analysis of multiple differentiated populations, all of which are common sources of variability within or between studies. We explore these questions with a pair of genome-wide datasets of the sort frequently used for assessing variation in wild populations. We focus initially on a ddRADseq dataset from one population of a common species and then consider a DArTseq dataset from a threatened species that covers multiple populations. We make some recommendations for assessing heterozygosity when study aims include comparisons of genetic variability across populations and with other studies.

## Materials and Methods

### Sequence data from the same population

We start by considering a single, well-mixed population. We use double digest restrictionsite associated (ddRAD) sequence data obtained from 100 female *Aedes aegypti* mosquitoes sampled from a 0.125 km^2^ area of Kuala Lumpur, Malaysia (Jasper et al., 2019). Note that as this ddRADseq dataset contains only females, and as *Ae. aegypti* mosquitoes do not have definable sex chromosomes but rather a small sex-determining region (Fontaine et al., 2017), we did not need to filter out genotypes at sex chromosomes.

We took subsamples from this population as follows.

#### Ten subsamples

We tested the effect of five population sample sizes (n = 10, 5, 4, 3, 2) on heterozygosity estimates by subsampling the 100 individuals, without replacement. We repeated the subsampling 10 times for each sample size n.

#### Nested subsamples

We tested the effect of six larger sample sizes (n = 50, 40, 30, 20, 10, 5) on heterozygosity estimates by subsampling the 100 individuals twice, without replacement. This can help indicate whether filtering choices produce similar patterns at large n as at small n. These subsamples were also used to test whether different filtering choices could produce variable heterozygosity estimates from the same sample of individuals. To reduce variation among subsamples of different size, we used a nested subsampling approach. The 100 individuals were randomly assigned to two groups, A and B, each of n = 50. Group A and Group B were then subsampled once at each n, but where each subsample could only include individuals that were included at the next highest n. For instance, the subsample at n = 30 could only contain individuals that were present in the n = 40 subsample to allow for a direct comparison between sample sizes.

### Sequence data from multiple populations

We considered the issue of multiple populations being included in a comparison by reanalysing a set of four populations of *Keyacris scurra* (Key’s Matchstick Grasshopper) taken from a larger set of sequencing data derived from a Diversity Arrays Technology (DArT) approach. *Keyacris scurra* has recently been listed as endangered and is currently restricted in range to refugia in south-eastern Australia. These four populations have experienced very low gene flow and are highly differentiated (pairwise *F*_ST_ = 0.14–0.28). The four populations were processed and sequenced together as part of the same project (Hoffmann et al., 2020). Note that no reference assembly is available for this species so the term “autosomal heterozygosity” here will also include sequence data from any differentiated sex chromosomes (or regions of chromosomes).

### Sequence processing

For the ddRADseq dataset, aligned sequences were built into a Stacks.v2 (Catchen et al., 2013) catalog with the program ref_map. For the DArTSeq dataset, sequence data were built into a *de novo* Stacks catalog using the program denovo_map, allowing for up to four mismatches within and between individuals. We analysed both datasets with the Stacks program “Populations”, which was used to estimate observed and expected heterozygosity for a range of filtering settings described below.

## Results

### Estimates based on polymorphic sites (SNP heterozygosity)

Our first aim was to see how a dataset filtered with settings typically used for assessing genetic structure (i.e. variation between populations) might perform when used to estimate heterozygosity (i.e. variation within populations). Analysis of genetic structure will usually consider only polymorphic sites (SNPs). When filtering SNPs, a common approach is to combine the entire data set, remove sites not genotyped in a sufficient number of individuals (typically 70-95%), and then filter out sites with a minor allele frequency (MAF) or minor allele count (MAC) that is not met globally (Lemopoulos et al., 2019; Mathur et al., 2019; Mulvena et al., 2020). Simulations suggest that a MAC ≥ 3 may be optimal for detecting population structure, as excluding rare alleles can lead to erroneous inferences of admixture but including singletons and doubletons can confound model-based inferences of structure (Linck & Battey, 2019).

#### Population comparisons using polymorphic sites only: effects of sample size

We start with a simple comparison of how global (N) and local (n) sample size affects SNP heterozygosity estimates. For this we use the ten subsamples (n = 10, 5, 4, 3, 2), which we analyse first individually (i.e. with each subsample run in a separate Stacks run) and then together (i.e. where the ten subsamples are run in a single Stacks run). To investigate these effects at n ≥ 5, we use the nested subsamples from Groups A and B, first analysing each subsample from Group A in individual runs, then analysing each pair of subsamples of equal n from A and B together. Filtering followed a standard approach for assessing genetic structure, retaining a single SNP from each RAD locus (--write-single-snp) which had no more than 20% missing data and that had a MAC ≥ 3. Observed and expected heterozygosities were estimated from these filtered polymorphic sites.

SNP heterozygosity estimates are shown in Fig. 1a-h and indicate how this type of filtering approach presents problems for comparing estimates across studies. In all cases, observed and expected heterozygosities were larger when fewer samples were used for estimation. Specifically, heterozygosities are biased by global N (total sample size in the analysis) rather than local n (sample size of each population), as evident from comparisons of subsamples of equal n analysed either in individual runs or together. Although these effects reduce as N increases, they persist even with n = 40 and N ≥ 80.

**Fig. 1:**
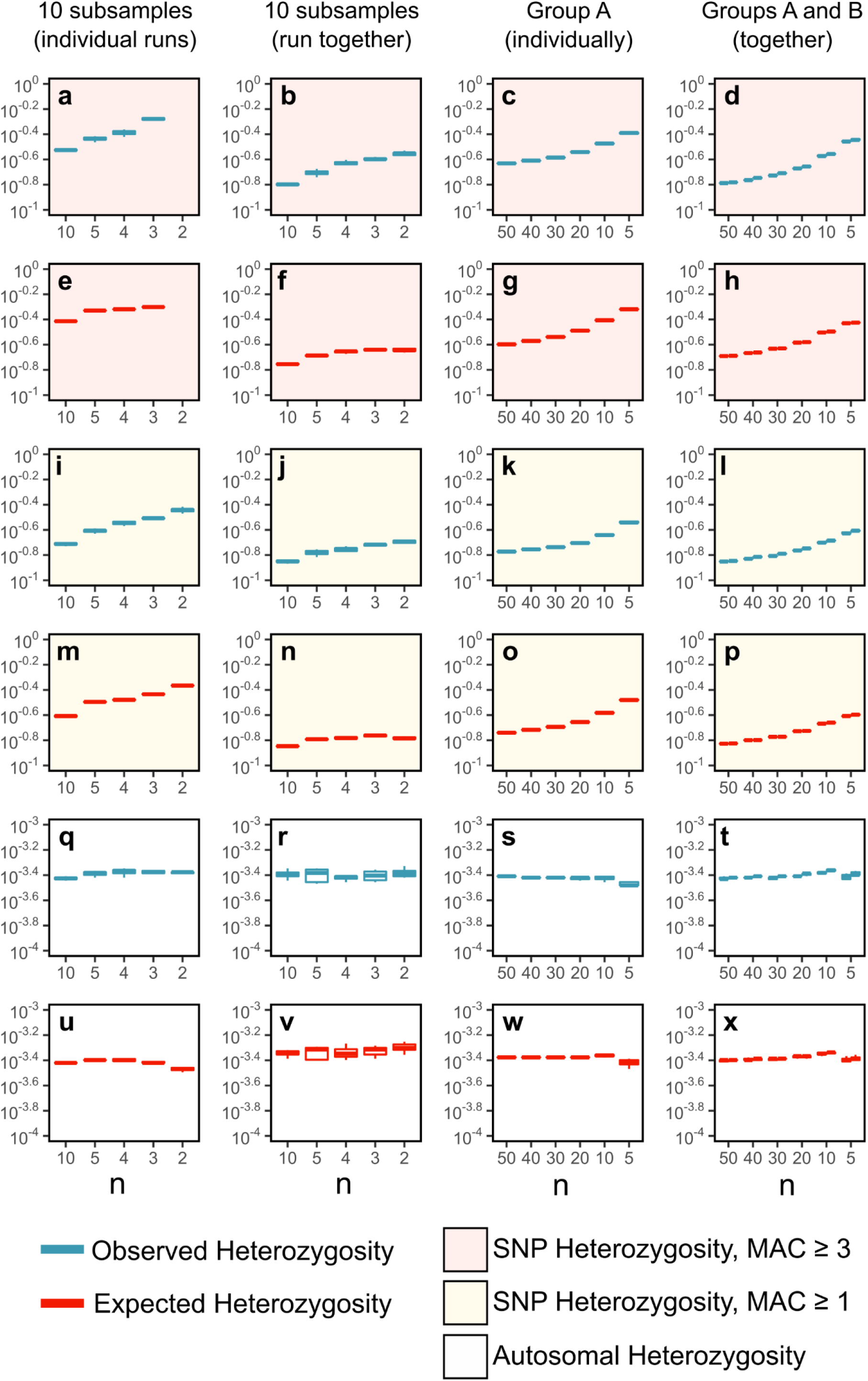
Boxplots showing effects of local and global sample size on heterozygosity estimates. Observed (blue; a-d,i-l,q-t) and expected (red; e-h,m-p,u-x) heterozygosities have been derived from three filtering treatments: polymorphic sites only, ≤ 20% missing data, MAC ≥ 3 (a-h); polymorphic sites only, 0% missing data, MAC ≥ 1 (i-p); polymorphic and monomorphic sites, 0% missing data (q-x). Treatments have been applied to four *Ae. aegypti* datasets described in the main text: ten subsamples each of size n, analysed in individual runs (a,e,I,m,q,u); ten subsamples each of size n, analysed together in a single run (b,f,j,n,r,v); single nested subsamples from Group A, each undergoing jack-knife resampling (c,g,k,o,s,w); nested subsamples from Group A and Group B analysed together, each undergoing jack-knife resampling (d,h,l,p,t,x). All Y-axes use a log-10 scale.

The source of this issue is that heterozygosity is generally lower for SNPs with rare alleles (where most individuals are homozygous for the common allele) than for SNPs with common alleles. For instance, for the n = 3 subsamples analysed in individual runs, all SNPs have minor alleles at 0.5 frequency when MAC ≥ 3 is applied (Fig. 1a, e), leading to expected heterozygosity of 0.5. As additional samples are added, SNPs with rare alleles become more likely to be detected, leading to lower heterozygosity estimates (c.f. Fig. S1). As MAC filtering is applied globally, heterozygosity is lower when populations are analysed together (Fig. 1b, f) as global sample size is ten times larger in these runs. However, even in these runs there were clear differences between SNP heterozygosity estimates for n = 10 (N = 100) and n ≤ 5 (N ≤ 50).

Considering these inconsistencies in SNP heterozygosity estimates when filtering datasets with ‘typical’ settings for genetic structure, we reran the above analyses with MAC ≥ 1 and selecting all SNPs rather than one per RAD locus. These analyses thus considered variation at all polymorphic sites including those with singletons and doubletons. We used a maximum missing data threshold of 0% (or as specified), to avoid potential artefacts caused by including sites with missing data (see Fig. 2). Including all polymorphic sites reduced the bias from sample size; however, similar patterns of strong bias were still observed (Fig. 1i-p). Thus, while including singletons and doubletons reduces sample size biases because rare alleles are then more likely to be detected in small samples, the biases will nevertheless persist when sites are filtered on the basis of polymorphism.

**Fig. 2:**
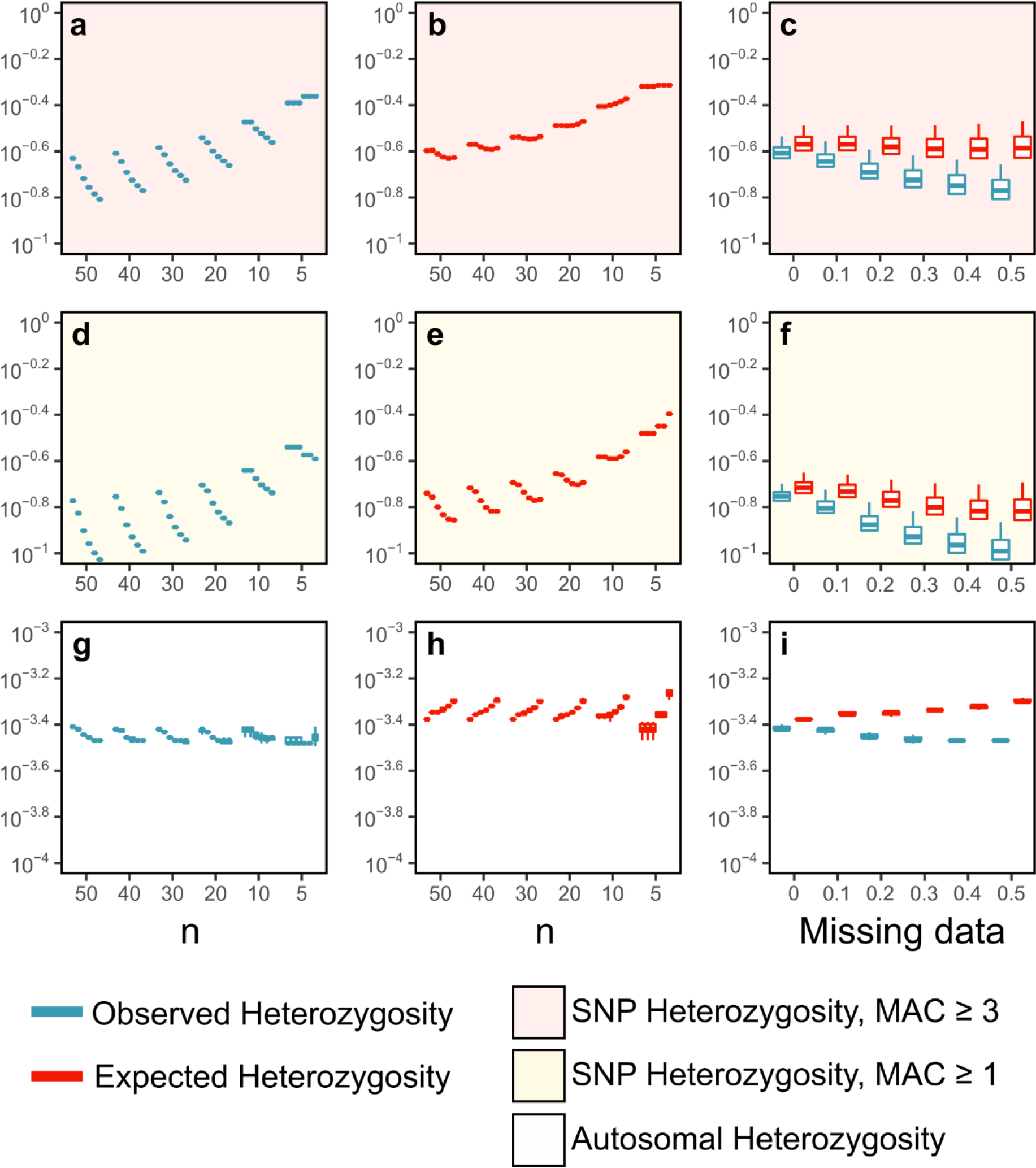
Boxplots showing effects of missing data thresholds on heterozygosity estimates. Observed (blue; a,d,g) and expected (red; b,e,h) heterozygosities have been derived from the nested subsamples from Group A following three filtering treatments: polymorphic sites only, MAC ≥ 3 (a-c); polymorphic sites only, MAC ≥ 1 (d-f); polymorphic and monomorphic sites (g-i). For each n, the subsample has been filtered using a progression of missing data thresholds: from left to right, 0%, 10%, 20%, 30%, 40%, 50%. Each estimate has undergone jack-knife resampling. Subfigures c,f,i aggregate results across all subsamples to show how observed (left) and expected (right) heterozygosities diverge with less stringent missing data thresholds. All Y-axes use a log-10 scale.

#### Population comparisons using polymorphic sites only: missing data thresholds

We investigated effects of missing data thresholds on SNP heterozygosity using the nested subsamples from Group A, filtered with thresholds of 0% (i.e. no missing data allowed), 10%, 20%, 30%, 40%, or 50%. Thus in each case variation in heterozygosity was assessed in a single population of size n (n = 50, 40, 30, 20, 10, 5). We compared results from the two filtering protocols described previously: a standard protocol for assessing genetic structure (one SNP per RADtag with MAC ≥ 3) and one that retains all polymorphic sites (MAC ≥ 1).

We see a considerable effect of missing data thresholds on SNP heterozygosity (Fig. 2a-f). Samples of larger n were more strongly affected by choice of missing data threshold, with stringent filtering tending to produce higher estimates. When 50 individuals were used with MAC ≥ 3 filtering (Fig. 2a), a 10% missing data threshold (a common parameter setting) produced an estimate for observed heterozygosity 1.22 times higher than filtering with a 30% threshold (also a common parameter setting). This effect was stronger with MAC ≥ 1 filtering (1.36 times higher; Fig. 2d). Expected heterozygosities were less biased by missing data thresholds but effects were still evident (Fig. 2b,e).

The higher observed heterozygosity estimates at 0% versus 50% thresholds might be expected if there is a correlation between errors and the presence of missing data at a site. In that case, errors at monomorphic sites with high missing data could be read as low frequency polymorphisms, pushing down heterozygosity estimates when N is large. Also, when singletons and doubletons are included more low-frequency errors would be included in calculations, leading to the stronger effects seen when MAC ≥ 1 filtering was applied.

Finally, the large population sizes in *Ae. aegypti* indicate this Malaysian sample is unlikely to contain inbred individuals, and thus we do not expect observed and expected heterozygosities to differ substantially. We note that when estimates of observed and expected heterozygosity are compared, these parameters are most similar when filtering with a 0% missing data threshold and start to diverge as this threshold is increased. These divergences are consistent across filtering types, from ~1.10 at 0% missing data to ~1.27 at 20% and ~1.50 at 50%. Less stringent missing data thresholds may thus introduce artefacts of differential observed and expected heterozygosities, which may lead to incorrect inferences of local breeding patterns.

In light of these inconsistencies, SNP heterozygosity appears prone to bias, regardless of whether filtering follows a typical protocol used for genetic structure or when considering every polymorphic site. This bias is demonstrated in Fig. S1, which shows how, when calculating SNP heterozygosity, the numerator remains proportionate to the number of heterozygous sites regardless of sample size, but the denominator is consistently biased downwards. This downward bias should diminish for very large N but for rare populations or small budgets this cannot be solved by sequencing more individuals. This limits the potential for SNP heterozygosity estimates in one study to inform the results of other studies.

### Estimates based on all polymorphic and monomorphic sites (autosomal heterozygosity)

Given the above challenges, we next explored how sample size and missing data affect autosomal heterozygosity, which considers both monomorphic and polymorphic nucleotides. We ran analyses on identical datasets to those used previously. For filtering, we used no MAC cut-off, and estimated heterozygosity across every site rather than every polymorphic site. In the output from the Stacks.v2 program “Populations”, this corresponds to the entries in the “# All positions (variant and fixed)” subsection. We used a maximum missing data threshold of 0% (unless specified differently).

#### Population comparisons using all sites: effects of sample size

When considering variation at all nucleotide sites, observed and expected heterozygosity estimates are far less affected by N than SNP heterozygosity estimates (Fig. 1q-x). Though there was some variability among subsamples of smaller n, observed heterozygosities of ~0.00039 and expected heterozygosities of ~0.00040 were consistently recorded. The similar estimates for these two parameters match expectations for this sample of Malaysian *Ae. aegypti*, where inbreeding is unlikely given the large size of mosquito populations and the spatial distribution of sampling.

These consistent estimates are expected when all sites are taken into account because for smaller samples the higher frequency of heterozygotes at polymorphic sites will be offset by the lower number of polymorphic sites overall. Heterozygosity estimates from a set of individuals thus correlate directly with population heterozygosity because sites are not first filtered by polymorphism (Fig. S1). In this sense, autosomal heterozygosity is a parameter that is both more robust to variation in study design and also a more accurate measure of genetic variation which can be used in comparisons across studies and organisms (Westbury et al., 2019).

#### Population comparisons using all sites: effects of missing data thresholds

Missing data thresholds had a smaller effect on autosomal heterozygosity than on SNP heterozygosity (Fig. 2g,h). Nevertheless, the same problematic pattern is clear in the divergence between observed and expected heterozygosities when sites with missing data are included, which was of equivalent magnitude to divergences in SNP heterozygosity (Fig. 2 i). Considering these results, we propose that heterozygosity estimation should exclude nucleotide sites that have any missing genotypes, as these may be more likely to contain errors or otherwise skew parameter estimates. While this filtering might at first appear overly stringent, autosomal heterozygosity is calculated using far more sites than SNP heterozygosity, and should normally be based on sufficient sites even after strict filtering. For example, when a 20% missing data threshold is used to estimate SNP heterozygosity (MAC ≥ 1) in 50 individuals, heterozygosity is estimated from 95,293 polymorphic sites. When a 0% missing data threshold is used to estimate autosomal heterozygosity (MAC ≥ 0), heterozygosity is estimated from 7,813,360 sites, of which 17,968 are polymorphic. This does not imply that the consistency in autosomal heterozygosity estimates is due to a larger number of sites; an increase in the number of sites will not resolve biases in SNP heterozygosity which reflect the sample size of individuals rather than sites (Fig. S1). Similarly, the specific number of nucleotides used in autosomal heterozygosity calculations may vary across studies, but should accurately reflect the degree of variation across the genome.

### Multiple population considerations

We first estimated heterozygosity for the four *K. scurra* populations using equal sample sizes of 10. Fig. S2 compares results of a 0% missing data threshold against a 20% threshold, showing that at 20%, the ratio of observed to expected heterozygosity is higher 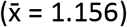 than at 0% 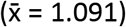, supporting our previous findings that missing data may bias the ratio of heterozygosities. Accordingly, we used a 0% missing data threshold in the following analyses.

#### Population comparisons: local sample size variation

Although global sample size analysed above had little effect on autosomal heterozygosity when n ≥ 10, we have yet to consider differences in sample size among populations. We estimated heterozygosity for the four *K. scurra* populations with one population (Goulburn) set at either half (5,10,10,10) or double (10,5,5,5) the size of the other populations compared to an equal population size. We compared results for autosomal heterozygosity and SNP heterozygosity following previous filtering settings (MAC ≥ 3 and MAC ≥ 1).

When 10 individuals are analysed from each population, the Goulburn, Hall and Wallendbeen populations all have similar heterozygosities, while Cooma is much lower (Fig. 3). There was no strong effect from unequal sample size for either autosomal heterozygosity (Fig. 3 e,f) or for SNP heterozygosity using all polymorphic sites (Fig. 3c,d), but filtering at MAC ≥ 3 revealed such an effect (Fig. 3a,b). The Goulburn population had either higher or lower heterozygosity than the Hall and Wallendbeen populations, depending on whether Goulburn had a greater or smaller n. This bias could lead to misinterpretation of relative SNP heterozygosities within studies when populations with different sample sizes are analysed together.

**Fig. 3:**
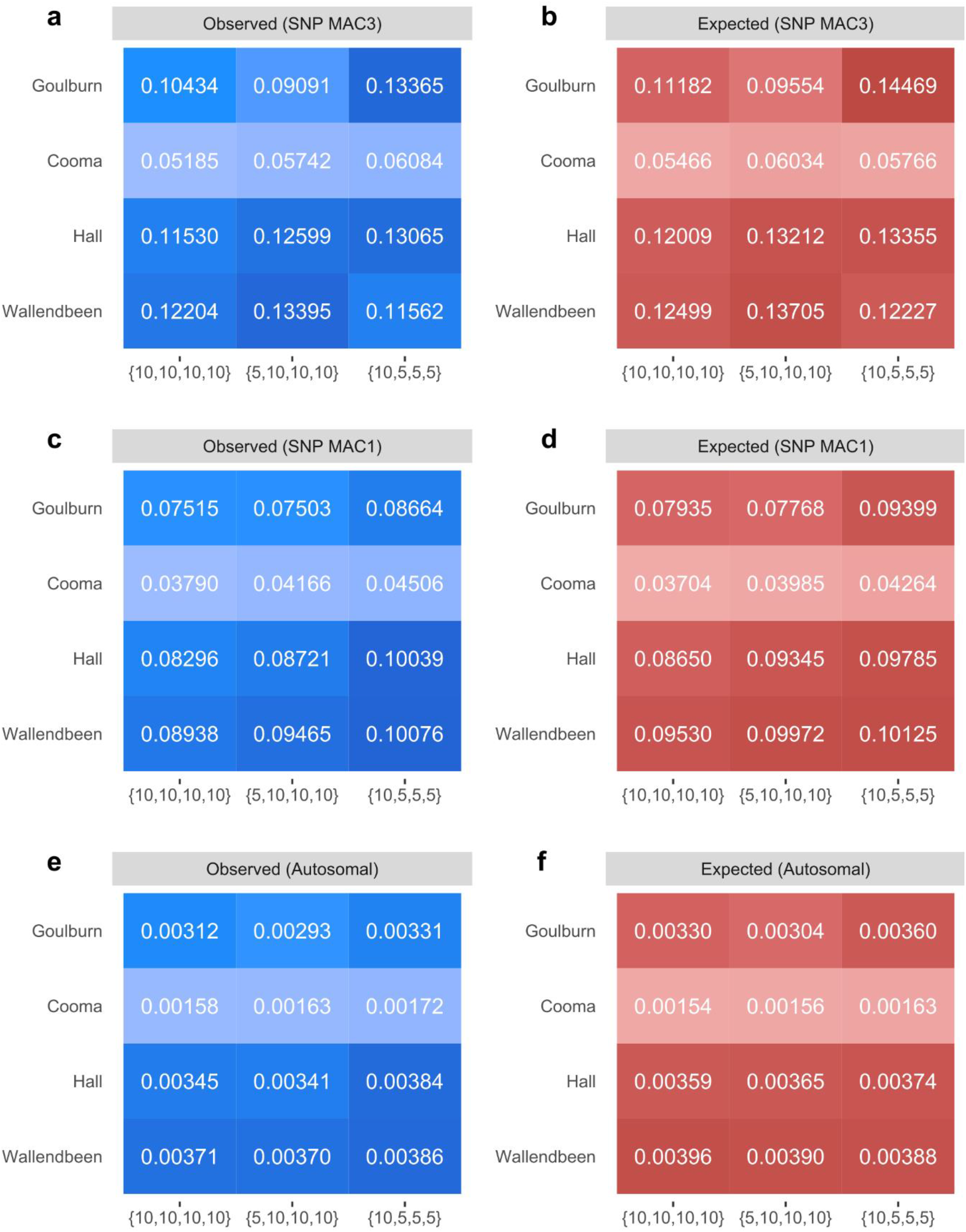
Effects of differential local sample size on heterozygosity estimates. Observed (blue; a,c,e) and expected (red; b,d,f) heterozygosities have undergone three filtering treatments: polymorphic sites only, MAC ≥ 3 (a,b); polymorphic sites only, MAC ≥ 1 (c,d); polymorphic and monomorphic sites (e,f). Numbers in brackets indicate sample sizes for the four *K. scurra* populations sequentially (Goulburn to Wallendbeen). Shading reflects similarity among numbers.

#### Population comparisons: impact of population structure

Next, we investigate the effects of combining genetically differentiated populations in an analysis. In terms of mtDNA variation and DArT SNPs, Cooma was separate from the other populations and particularly Wallendbeen (Hoffmann et al., 2020). We estimated SNP and autosomal heterozygosities for each of the four *K. scurra* populations analysed individually (i.e. with each population in a separate Stacks run) and compared this to populations analysed together (i.e. with all populations in a single Stacks run).

Autosomal heterozygosity is unaffected by whether differentiated populations are analysed individually or together (Fig. 4b). However, strong biases on SNP heterozygosity are evident (Fig. 4a). When the four populations are analysed individually, estimates are much higher than when analysed together. Additionally, the population at Cooma, which otherwise recorded the lowest heterozygosity of the four populations, has higher heterozygosity than all other populations when analysed by itself.

**Fig. 4:**
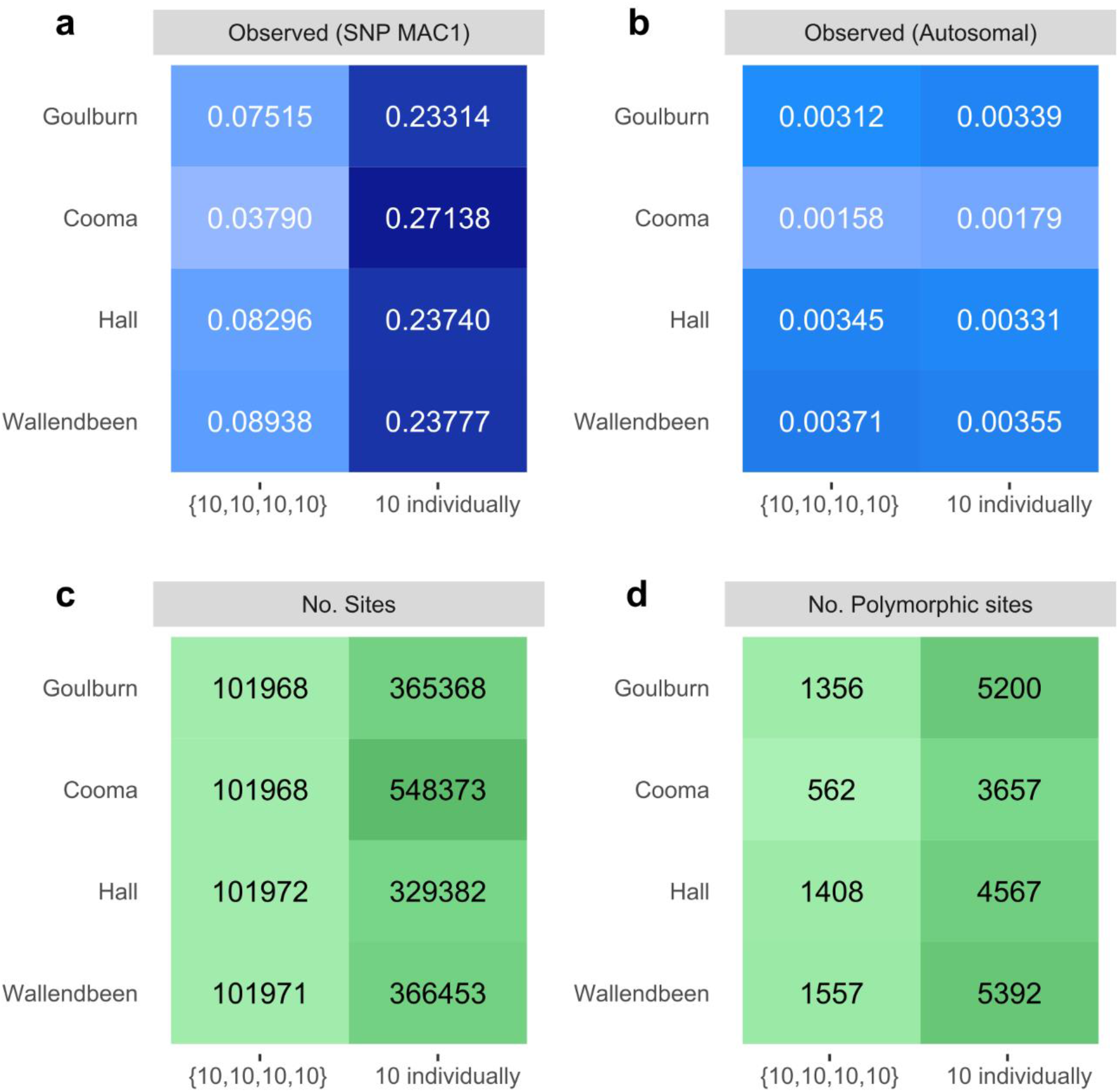
Effects of population genetic structure on heterozygosity estimates. Observed SNP heterozygosities (a) and autosomal heterozygosities (b) are presented for the four *K. scurra* populations which have either been analysed together in a single run or in separate runs individually. The number of sites (c) and number of locally polymorphic sites (d) retained after filtering are also presented. Shading reflects similarity among numbers.

As allele frequencies vary among these populations, many variant sites will only be polymorphic in one or two populations and are monomorphic in the others. Thus when populations are analysed together, estimates for each population will include these variant sites that are locally monomorphic, leading to lower heterozygosity estimates than when analysed individually.

A similar explanation accounts for the sharp variation in estimates at Cooma. When analysed individually, Cooma recorded fewer polymorphic sites (3235) than the other populations (4693, 4734, 5182); this pattern was also observed when populations were analysed together. However, heterozygosity at these 3235 polymorphic sites was higher than at the other populations.

These findings show how SNP heterozygosity estimates represent different parameters when populations are analysed in individual runs compared to when they are analysed alongside other populations. When analysed individually, the SNP heterozygosity of a population is equal to autosomal heterozygosity multiplied by the proportion of sites that are polymorphic (Fig. S1). When analysed with other populations, the SNP heterozygosity of a population will be shaped by whichever other populations are included, as the structure between these populations will determine which locally monomorphic sites are called as SNPs. Accordingly, SNP heterozygosities, if they are to be reported, should probably be calculated from populations analysed individually, and the total number of polymorphic sites should also be reported for each population to provide further context to the heterozygosity estimates. For autosomal heterozygosity, analysing multiple populations at once introduces no biases while conferring no advantage, but does reduce the number of retained sites (Fig. 4 c,d). It follows that calculations of autosomal heterozygosity should analyse each population in individual runs. For observed autosomal heterozygosity, this could be extended to analysing each individual in turn if needed.

### Comparing heterozygosity estimates

A final consideration concerns how to interpret heterozygosity estimates across studies. We have proposed several guidelines for filtering data to allow cross-study comparisons. The most important of these is that heterozygosity estimates should be derived from variation at both monomorphic and polymorphic sites. Table 1 compares variation in SNP heterozygosity with variation in autosomal heterozygosity for the four *K. scurra* populations analysed individually. We did not compare populations when analysed together due to the confoundment of SNP heterozygosities in these analyses (Fig. 4).

**Table 1.** Magnitude of heterozygosity differences between populations for SNP and autosomal heterozygosity. Calculations are based on the results from Fig. 4. MAX/MIN is the ratio between the largest score and the smallest score.

| | SNP heterozygosity (MAC ≥ 1) | | Autosomal heterozygosity | |
| --- | --- | --- | --- | --- |
|  | Observed | Expected | Observed | Expected |
| MAX/MIN | 1.164 | 1.064 | 1.983 | 2.230 |
| Coefficient of variance | 0.063 | 0.024 | 0.236 | 0.267 |

For *K. scurra*, variation in autosomal heterozygosity is approximately twice as large as variation in SNP heterozygosity (Table 1). A large difference in SNP heterozygosity might not be detected even when comparing populations with very low and very high levels of genetic variation because the exclusion of monomorphic sites in each population will reduce differences in genetic variability among the populations.

## Discussion

Comparisons of heterozygosity across populations and species are frequently used to inform management decisions in conservation programs. An example of a relevant management action involves selecting populations for genetic and evolutionary rescue, which aims to decrease levels of inbreeding and increase levels of genetic variation in target populations through targeted introductions of individuals from other populations (Hoffmann et al., 2020; Whiteley et al., 2015). The usefulness of source populations for conservation translocations is to a large extent determined by their genetic variability (Ørsted et al., 2019; Reid et al., 2016). Tracking changes in heterozygosity across time can also be a worthwhile means of tracking the genetic health of threatened populations (Mitrovski et al., 2008) and is particularly useful for determining outcomes of management interventions (Weeks et al., 2017). All of these objectives require that heterozygosity estimates are comparable across populations within a study and across different studies.

In this paper, we have shown that filtering genome-wide sequence data using optimal settings for detecting genetic structure will produce heterozygosity estimates that are poorly-suited to these comparisons. Specifically we show that heterozygosity estimates that consider only polymorphic sites (SNP heterozygosity) are always biased by global sample size (N), with smaller sample sizes producing larger heterozygosity estimates.

Heterozygosity estimates that consider monomorphic and polymorphic sites (autosomal heterozygosity) do not suffer from these biases. We also found that when sites with missing data are included, observed and expected heterozygosity estimates diverge, with the divergence proportional to the amount of missing data permitted. When multiple populations were analysed together, SNP heterozygosity estimates were additionally biased by allele frequency differences among populations. While analysing populations together did not bias autosomal heterozygosity, it conferred no advantages over analysing populations individually but reduced the number of available sites due to missing data filtering.

Following this, we propose three general guidelines that should help meaningful comparisons: (i) studies aiming to summarise population genetic variation should report autosomal heterozygosity, either by itself or alongside SNP heterozygosity; (ii) sites with missing data should be omitted from heterozygosity calculations; and (iii) populations should be analysed in independent runs. Although we have not explicitly investigated the importance of sequencing coverage, this is widely known to be critical for accurately identifying heterozygotes (Nielsen et al., 2011). This being the case, our findings that heterozygosity estimates can be consistent even at low n (Fig. 7a) point to the optimal design for heterozygosity being deep sequencing of a small number of individuals (perhaps 5-10) from each population, rather than shallower sequencing of many individuals. Previous work has found these numbers to be adequate (Nazareno, Bemmels, Dick, & Lohmann, 2017).

SNP heterozygosity is frequently the only measure of heterozygosity reported (Bock et al., 2018; Chen et al., 2016; Jones et al., 2012; Mathur et al., 2019; Surbakti et al., 2020). Although there may be cases where SNP heterozygosity is the appropriate parameter, it will be subject to biases from sample size that do not affect autosomal heterozygosity, making the latter a better ‘default’ choice for reporting variation in populations. As taxa of conservation interest are frequently rare or difficult to sample in large numbers, large sample sizes for all populations would be difficult to achieve. SNP heterozygosity also does not capture all the variation in the genome. For instance, *K. scurra* from Cooma had a higher SNP heterozygosity estimate (when calculated in an independent run) yet fewer polymorphic sites than the other populations; this low level of polymorphism was evident in the autosomal heterozygosity estimate.

While we advocate autosomal heterozygosity as a default choice for reporting genome-wide averages, there may be circumstances where variation at a specific set of sites is of interest (e.g. Chen et al., 2016). In this case, heterozygosity can be estimated free of bias provided that this specific set of sites is not further filtered by polymorphism when new populations are analysed, and thus sites should be retained even if the new population is locally monomorphic. Returning to the work of Nazareno et al. (2017), the consistency of their heterozygosity estimates across subsamples of different size is due to these subsamples not being refiltered by polymorphism. While Nazareno et al. (2017) effectively demonstrates how heterozygosity can be estimated with small samples, this approach would not be applicable to comparisons across populations. Likewise, assessing variation at a specific set of sites may evade sample size biases, but other biases may be introduced when assessing variation in populations with different allele frequencies to those in populations used to select the initial set of variable sites.

Whole-genome sequencing studies frequently report autosomal rather than SNP heterozygosity (Gopalakrishnan et al., 2017; Westbury et al., 2019). However, this methodology has been less commonly applied in reduced-representation sequencing studies such as those using RADseq or DArTseq markers. One reason for this may be that whole-genome studies frequently analyse single individuals rather than a sample, and as SNP heterozygosity calculated from a single diploid individual will always be equal to 1 (Fig. S1) this approach would not be considered. Another reason may be that reduced-representation protocols are commonly applied to wild populations of understudied organisms, where the first step is typically to analyse genetic structure among populations. Monomorphic sites are uninformative for most analyses of genetic structure and thus these are typically removed during filtering, and this remaining set of SNPs will be used to estimate both genetic structure and heterozygosity. Here we propose that analysis of genetic structure (variation between populations) and heterozygosity (variation within populations) be treated as distinct targets and filtered using different parameter settings.

When expected heterozygosity is higher than observed heterozygosity, it is often treated as evidence for local inbreeding (Hoffmann et al., 2020), an important parameter that can indicate a need for genetic intervention in threatened species (Ralls et al., 2018). However, Fig. 2 shows that inferences of inbreeding in the presence of missing data may be confounded, as observed and expected heterozygosities quickly diverge as sites with missing data are included, leading to F_IS_ > 0. These opposing effects of missing data filters on observed and expected heterozygosities are consistent with sequencing error rates being higher at nucleotide positions where there are some missing genotypes. A possible explanation is that sequencing errors are more likely at monomorphic sites; these sites are more common, and more likely to be retained than errors at polymorphic sites which may introduce a third allele and be removed from the dataset (most filtering protocols retain only biallelic sites). Monomorphic sites with errors may therefore be coded as low-frequency SNPs. Including them could affect the observed and expected autosomal heterozygosity estimates differently depending on whether errors are mostly coded as homozygous or heterozygous.

Despite these issues, the *K. scurra* data also indicate clear instances where inbreeding levels differ across populations. We note that for *K. scurra* from Cooma, observed and expected heterozygosities were almost identical, while expected heterozygosities are 8-15% higher in the other populations, suggesting a situation where inbreeding in Cooma is low but genetic variability is also low. An accurate assessment of inbreeding in populations versus low genetic variation is important when making recommendations around genetic mixing of threatened populations, which can target both the masking of deleterious genes expressed as a consequence of inbreeding as well as problems from low levels of genetic variability (Hoffmann et al., 2020; Ralls et al., 2018; Weeks et al., 2011). It would be worth further evaluating optimal filtering strategies for assessing the ratio of observed and expected heterozygosities, using a set of populations where inbreeding is known to occur in some populations but not others.

This study has highlighted issues with estimating population heterozygosities from SNP data directly, and shown that autosomal heterozygosity estimates are more robust to the influence of sample size and are likely to be more comparable across studies. We provide general guidelines for estimating population heterozygosity from genome-wide sequence data that are usually different from guidelines for estimating other population genetic parameters such as gene flow, population structure, relatedness and effective population size. As in previous assessments (Linck & Battey, 2019), our results demonstrate that SNP datasets need to be carefully evaluated when they are used to obtain genetic parameters for populations that inform management decisions.

## Acknowledgements

This research was supported by the Australian Research Council (Discovery Grant DP190100990), the Wellcome Trust (Grant no. 108508) and the National Health and Medical Research Council (Program Grant no. 1132412; Fellowship Grant no. 1118640), and facilitated by use of the Nectar Research Cloud.

## Data Archiving

Aligned. bam files for 100 *Ae. aegypti* are available through the NCBI SRA, accession number PRJNA735025. Sequence data for *K. scurra* is available from https://www.ncbi.nlm.nih.gov/bioproject/702007.

## Author Contributions

AAH and ARW conceived the study; TLS, MJ, AAH and ARW designed the methodology; TLS and MJ analysed the data; TLS, AAH and ARW led the writing of the manuscript. All authors contributed critically to the drafts and gave final approval for publication.

## Supplementary Figures S1 – S2

**Fig S1.**
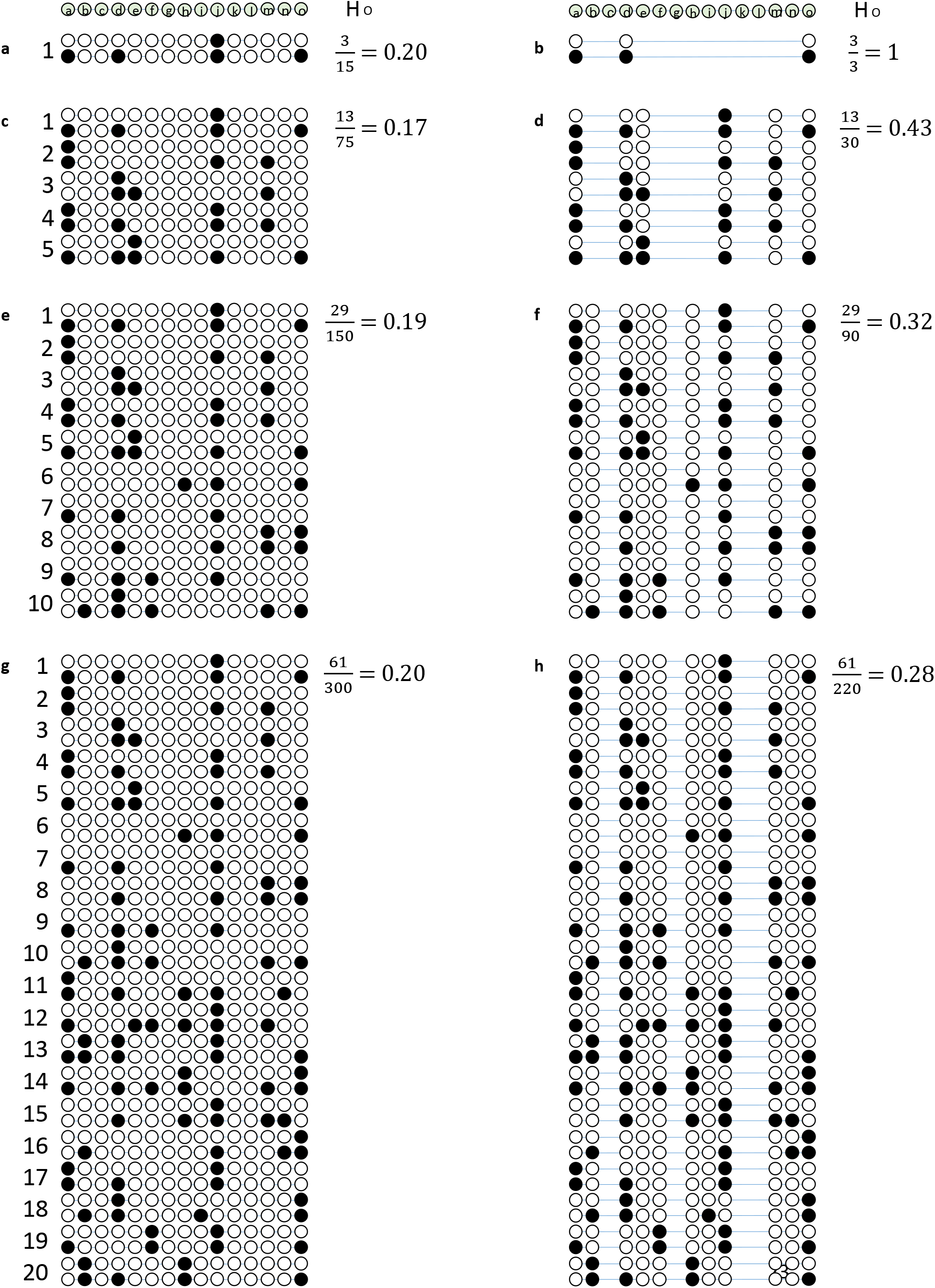
Estimating heterozygosity in a hypothetical population of diploid individuals. Analyses consider population samples of n = 1 (**a**,**b**), 5 (**c**,**d**), 10 (**e**,**f**) and 20 (**g**,**h**). Observed heterozygosity is calculated using all sites irrespective of polymorphism (autosomal heterozygosity: **a**,**c**,**e**,**g**) or using SNPs only (SNP heterozygosity: **b**,**d**,**f**,**h**). Individuals have been genotyped at 15 sites, and the circles indicate the unphased genotype at each site. When autosomal heterozygosity is calculated, the numerator and denominator remain proportionate to the number of sites evaluated (denominator) and the number of heterozygous sites (numerator) regardless of sample size. When SNP heterozygosity is calculated, the denominator is biased by whether SNPs with rare alleles are detected in the sample, which will remain an issue even at large n. These biases will be present in both observed and expected heterozygosity. Note that in actual sequence data the proportion of monomorphic sites will be much higher than shown here, producing smaller autosomal heterozygosity estimates.

**Fig S2.**
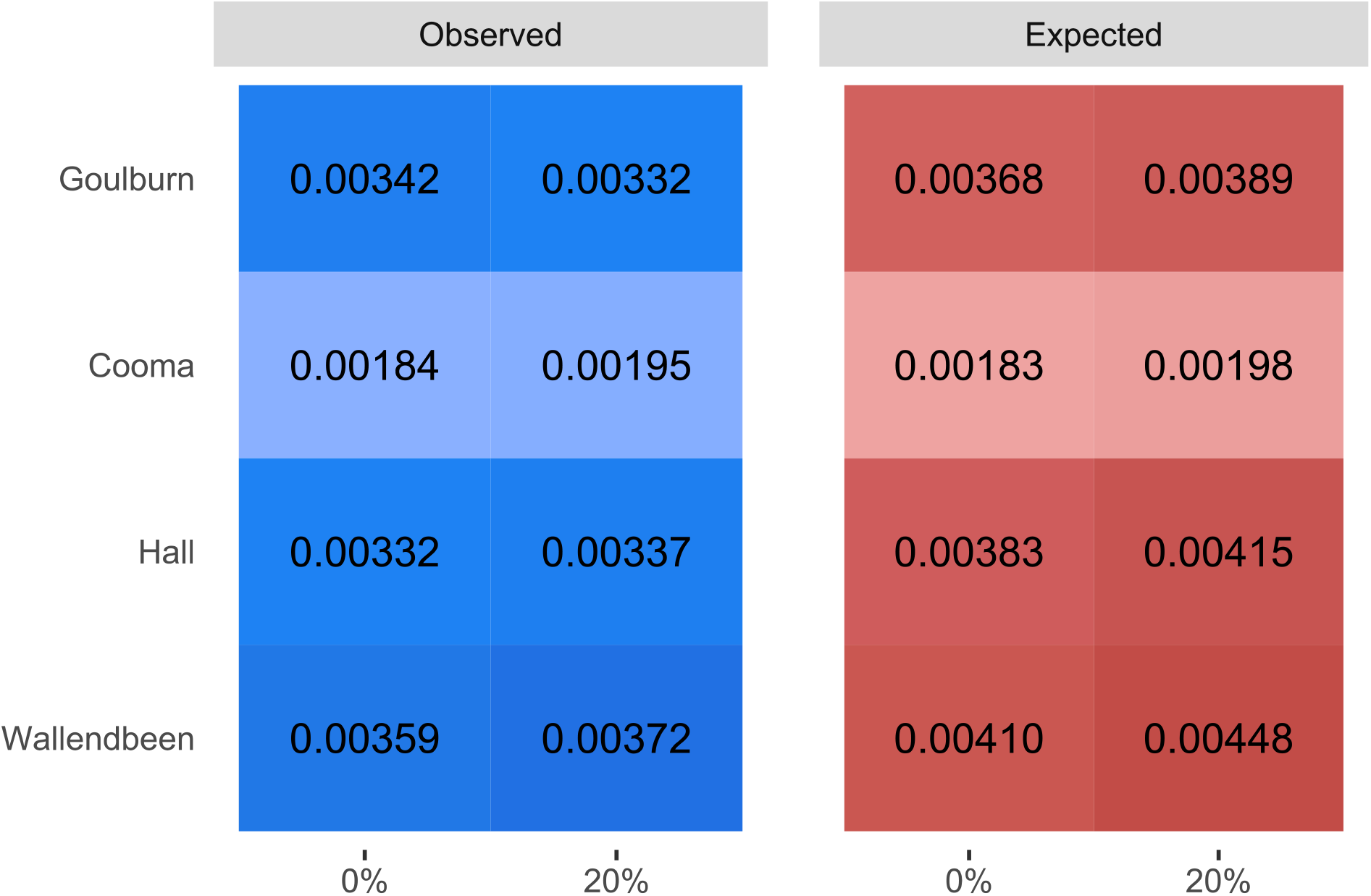
Estimates of observed and expected autosomal heterozygosity in four *K. scurra* populations filtered with 0% vs 20% missing data thresholds. The mean heterozygosity ratio between observed and expected values at 0% is 1.09, and at 20% it is 1.16. Entries with similar shading are more similar to each other.

